# A genetic investigation of sex bias in the prevalence of attention deficit hyperactivity disorder

**DOI:** 10.1101/154088

**Authors:** Joanna Martin, Raymond K. Walters, Ditte Demontis, Manuel Mattheisen, S. Hong Lee, Elise Robinson, Isabell Brikell, Laura Ghirardi, Henrik Larsson, Paul Lichtenstein, Nicholas Eriksson, Thomas Werge, Preben Bo Mortensen, Marianne Giørtz Pedersen, Ole Mors, Merete Nordentoft, David M. Hougaard, Jonas Bybjerg-Grauholm, Naomi Wray, Barbara Franke, Stephen V. Faraone, Michael C. O’Donovan, Anita Thapar, Anders D. Børglum, Benjamin M. Neale

## Abstract

Attention-deficit/hyperactivity disorder (ADHD) shows substantial heritability and is 2-7 times more common in males than females. We examined two putative genetic mechanisms underlying this sex bias: sex-specific heterogeneity and higher burden of risk in female cases. We analyzed genome-wide common variants from the Psychiatric Genomics Consortium and iPSYCH Project (20,183 cases, 35,191 controls) and Swedish population-registry data (N=77,905 cases, N=1,874,637 population controls). We find strong genetic correlation for ADHD across sex and no mean difference in polygenic burden across sex. In contrast, siblings of female probands are at an increased risk of ADHD, compared to siblings of male probands. The results also suggest that females with ADHD are at especially high risk of comorbid developmental conditions. Overall, this study supports a greater familial burden of risk in females with ADHD and some clinical and etiological heterogeneity. However, autosomal common variants largely do not explain the sex bias in ADHD prevalence.

## Introduction

Attention-deficit/hyperactivity disorder (ADHD) is a common (~5% childhood prevalence) and highly heritable (70-80%) neurodevelopmental disorder^1,2^. Recent large-scale genome-wide association studies (GWAS) suggest the involvement of thousands of genetic risk variants across the spectrum of allele frequencies^3–7^. The first robust genome-wide significant single nucleotide polymorphisms (SNPs) have recently been identified in a GWAS meta-analysis of 20,183 ADHD cases and 35,191 controls/pseudo-controls^7^. While such efforts are beginning to shed light on the biology of ADHD, secondary genome-wide analyses can address important issues regarding the etiological and clinical heterogeneity of ADHD.

One striking epidemiological observation about childhood ADHD is that males show a 2-7 times higher diagnosis rate compared to females^1,8^. The male excess is more pronounced in individuals ascertained from clinics than from the community, and this sex difference attenuates in adulthood^2^. The reasons for the substantial difference in prevalence in childhood are unclear. Here we present a series of analyses aimed at elucidating the basis for these differences.

### Sex-specific heterogeneity

One possibility is that ADHD in females is qualitatively different from ADHD in males. Although a review of twin studies found no difference in overall heritability estimates by sex in ADHD^9^, this does not necessarily imply that the same genetic risk variants are involved in the etiology of ADHD in males and females. If ADHD in clinically-diagnosed males is distinct from ADHD in diagnosed females, then this could yield differences in the prevalence. Sex-based genetic heterogeneity in common variants has been shown for several complex human traits (e.g. blood pressure and waist-hip ratio)^10,11^. Here, we used a genome-wide assessment of the genetic correlation of ADHD in males and females to determine, whether genetic heterogeneity from common variation contributes to the observed biased prevalence.

The absence of extensive sequencing data currently precludes analogous analyses of rare genetic variants. Instead, to evaluate whether such variants play differential roles in males and females with ADHD, we used risk for comorbid brain-related developmental disorders (i.e. ASD, intellectual disability, epilepsy, motor developmental delay) and rare syndromic phenotypes (i.e. congenital malformations, syndromes related to chromosomal abnormalities) as a proxy for possible presence of de novo or rare segregating alleles. Rare, highly deleterious (including non-inherited) genetic variation has been implicated in such phenotypes^12–22^. Indeed, comorbid ID in the context of ADHD has been associated with an increased likelihood of an individual being a carrier of a large, rare CNV^23^. It has also long been known that rare genetic syndromes (e.g. Fragile-X syndrome, velo-cardio-facial syndrome) are associated with ADHD^24,25^. Evidence for an increase in such comorbid conditions in females with ADHD, when compared to affected males, would imply a more severe, even ‘syndromal’, phenotypic presentation of ADHD in a higher proportion of females. Such more complex phenotypic presentations are arguably more likely to be linked to deleterious rare mutations. A higher rate of these comorbidities in females would also be consistent with clinical heterogeneity, which may be relevant to the differences observed in prevalence.

### Female protective effect

Aside from heterogeneity, the differences in prevalence may be a consequence of a ‘female protective effect’, whereby females are resilient to developing ADHD, and thus require a higher burden of genetic liability to develop the disorder. Three previous family studies have observed indirect evidence for this hypothesis, in the form of increased risk of ADHD in first-degree relatives (twins or parents) of affected females compared to affected males, suggesting that families with an affected female may have an overall higher burden of genetic risk^26–28^. Not all studies report an increase in the recurrence rate of ADHD in relatives of female probands however^29,30^. Two molecular genetic studies have more directly tested the female protective effect hypothesis using ADHD GWAS discovery data to calculate the burden of common risk alleles, as estimated by polygenic risk scores (PRS), in independent samples. In both studies, female children with ADHD-related phenotypes had higher PRS for ADHD than affected males^3,31^. Although these preliminary studies are consistent with the family studies mentioned above, they were based on small discovery GWAS. Additional tests utilizing large GWAS datasets are needed to test whether there is an increased burden of common genetic risk variants in females with ADHD.

We present a series of analyses to test the qualitative and quantitative difference hypotheses for the biased sex prevalence in ADHD. First, genome-wide common variant data from the autosomes were used to estimate the genetic correlation of ADHD in males and females. We then used population registry data to examine, whether females with ADHD are at an increased risk of comorbid developmental conditions compared to affected males. Next, genome-wide SNP data were used to test whether females diagnosed with ADHD carry a higher burden of common risk variants than affected males. Finally, we tested whether relatives of females with ADHD are at an increased risk of ADHD compared to relatives of diagnosed males.

## Results

### Sex-specific heterogeneity

#### Genetic correlation

Figure 1 displays genetic correlation (r_g_) results for male and female ADHD from bivariate analyses using both GREML and LDSC (see Table S1 for exact estimates). The LDSC r_g_ estimate in the full dataset (PGC+iPSYCH) was very high, near 1. Similar results were found for bivariate GREML analyses in both iPSYCH and PGC, as well as for the LDSC analyses in the iPSYCH dataset. The estimate using LDSC in the PGC dataset was lower, but large standard errors were seen in the PGC dataset for both methods.

**Figure 1.**
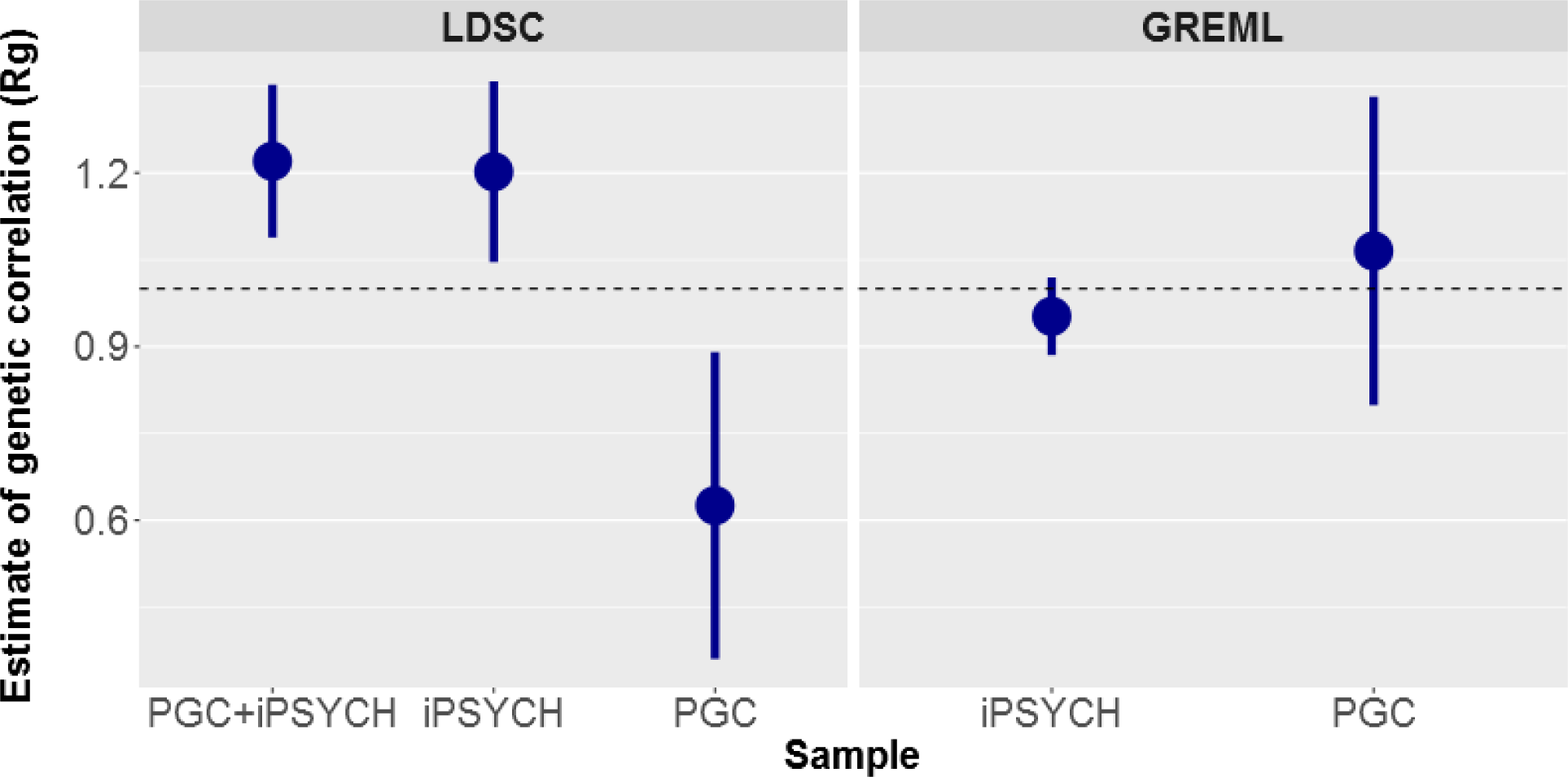
Genetic correlation estimates for ADHD in males and females obtained from GREML and LDSC for the iPSYCH, PGC, and combined PGC+iPSYCH datasets. Because of strict restrictions on raw individual genotype access and transfer, GREML analyses could only be performed separately in the PGC and iPSYCH samples. Bars display standard errors. The horizontal dashed line indicates a genetic correlation of 1.

Additional cross-dataset and cross-sex LDSC genetic correlation estimates were used to assess the extent of heterogeneity across the two different sub-samples (PGC and iPSYCH); see Figure S1 and Table S2. The genetic correlation between PGC and iPSYCH samples was not significantly different from 1 (r_g_(se)=1.13(0.22)). Cross-dataset r_g_ for PGC males with iPSYCH males and females were also not significantly different from 1. Estimates for PGC females with iPSYCH were lower (significantly different from zero and from 1), though this is likely to be at least partly related to the small sample size of the PGC females (N=1,067 cases and N=5,178 controls).

SNP-h^2^ estimates are displayed in Figure S2 and Table S1. SNP-h^2^ was estimated at 0.123 (se=0.025) in females and 0.247 (se=0.021) in males. Down-sampling male cases and controls randomly to match the female sample size showed more similar SNP-h^2^ estimates (See Figure S2 and Table S3). Results varying the relative population prevalence assumed are shown in Table S4. Figure 2 summarizes these results, illustrating the impact of the assumed male:female ratio (which affects assumed sex-specific population prevalence rates) and sample size on SNP-h^2^ estimates; SNP-h^2^ estimates increased in males and decreased in females as the ratio was increased and down-sampling male cases and controls gave similar SNP-h^2^ estimates in both sexes.

**Figure 2.**
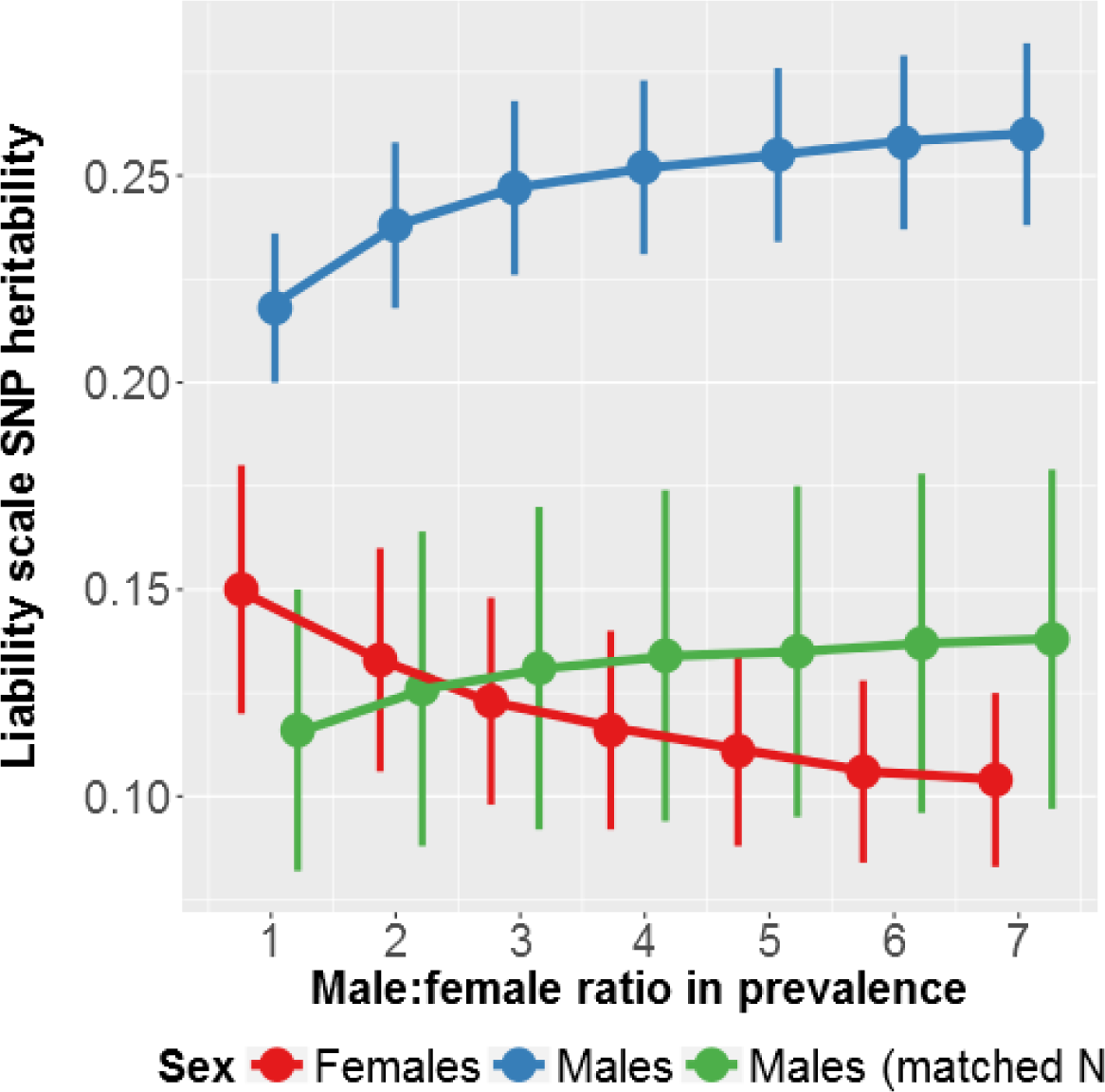
Sex-specific SNP-heritability estimates for PGC+iPSYCH using LDSC, varying the assumed population prevalence based on different male:female ratios (ranging from 1:1 to 7:1). Estimates are presented for the total available sample of males as well as for a down-sampled set of male cases and controls to match the available sample size in females.

#### GWAS

Sex-specific QQ and Manhattan plots are shown in Figure 3. There were 3 independent genome-wide significant loci in the male-only GWAS (N=14,154 cases & 17,948 controls). No SNPs surpassed the threshold for genome-wide significance in the female-only GWAS (N=4,945 cases & 16,246 controls). The top 10 LD-independent SNPs for each GWAS, annotated with the nearest gene are displayed in Table S5.

**Figure 3.**
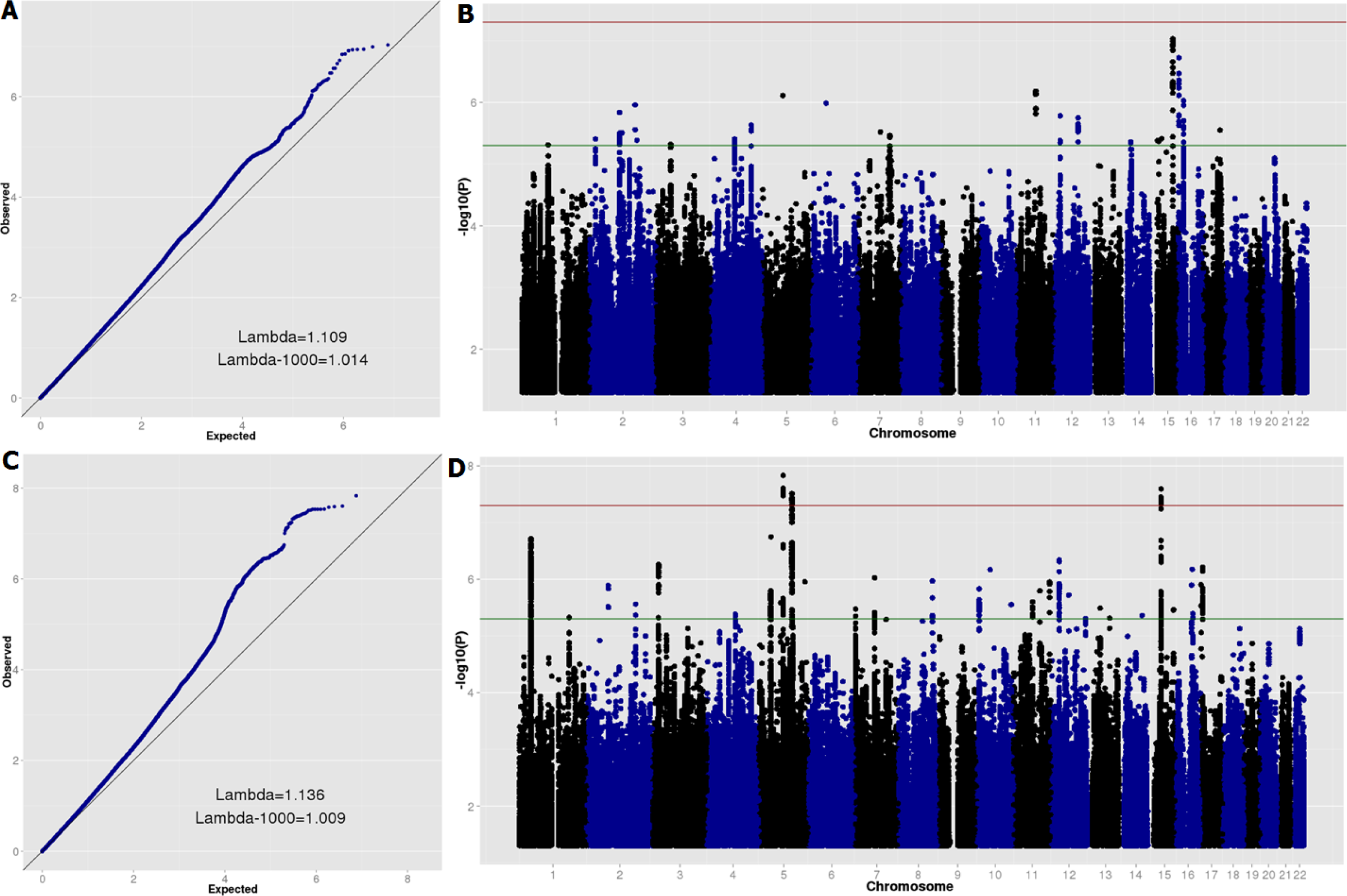
QQ and Manhattan plots for sex-specific genome-wide association meta-analyses. a) Female case-control analysis QQ plot; b) Female case-control analysis Manhattan plot; c) Male case-control analysis QQ plot; d) Male case-control analysis Manhattan plot. In figures b) and d) the horizontal red line indicates genome-wide significance (p<5E-8) and the horizontal green line indicates suggestive sub-threshold signals (p<5E-6).

Several secondary analyses support the high genetic correlation results, suggesting that there is little or no difference in the ADHD results for males and females. First, no genome-wide significant heterogeneity is observed when meta-analyzing the male and female GWAS results (see Figure S3). Second, a GWAS of sex-by-genotype interactions for ADHD identifies no individual variants with differential effects by sex, nor does it show any deviation from the null distribution of test statistics genome-wide (see Figure S4). Similarly, GWAS results for ADHD in the combined sample with or without including sex as a covariate are nearly perfectly correlated, with a low standard error (r_g_=0.97, se=0.007). Narrowing the GWAS focus to only ADHD cases also finds no genome-wide significant differences between male cases and female cases (Figure S5). Although some genome-wide inflation is observed for this final analysis in the iPSYCH sample, it is not replicated in the PGC data and appears to be attributable to association in only one locus driven by a single low-frequency genotyped SNP (MAF=0.02). Investigation of this locus shows no support for differences between male and female cases in neighboring genotyped SNPs, suggesting that the signal at this locus is likely a technical artefact (see Figure S6).

#### Epidemiological analyses

To test for heterogeneity in males and females diagnosed with ADHD, we examined the association between having an ADHD diagnosis and risk of having a comorbid developmental phenotype using epidemiological data from Sweden. In particular, we were interested in whether there is an interaction between sex and ADHD as it pertains to these comorbidities. Table S6 displays the frequency of the disorder categories examined as well as the proportion of individuals affected, overall and split by ADHD case status and sex. The male:female ratio for ADHD was 2:1 in the Swedish population. Analyses revealed that male and female ADHD cases are at a higher risk for all diagnostic categories, as compared with sex-matched population controls (see Table 1). Significant ADHD-by-sex interactions were observed for ASD and congenital malformations (CM), suggesting that, although in the context of ADHD both sexes are at an increased risk of comorbid ASD and CM, compared with controls, the increase in risk is even higher in females than in males. A nominally significant association was observed for the interaction term for ID as an outcome, which did not survive correction for multiple testing (Bonferroni correction for 6 independent tests: p-value threshold=0.0083). Secondary analyses of severity of ID (where this information was available) indicated that this weak association signal came from mild ID (IQ=50-70), not moderate (IQ=35-49) or severe/profound (IQ<35) ID (see Table S7). Interaction terms were non-significant for epilepsy, developmental coordination disorder, or chromosomal abnormalities.

**Table 1:** Results of logistic regression analyses of ADHD case status on comorbid developmental conditions, stratified by sex, in the Swedish population sample (Total N=1,952,542)

| Outcome | Sex | % of ADHD cases with outcome | Sex-specific association |  |  |  | ADHD by sex interaction |  |  |  |
| --- | --- | --- | --- | --- | --- | --- | --- | --- | --- | --- |
|  |  |  | OR | LCI | UCI | p | OR | LCI | UCI | p |
| ASD | Males | 14.82 | 18.91 | 18.33 | 19.50 | **<2.2E-308** | 1.52 | 1.44 | 1.61 | **2.9E-50** |
|  | Females | 12.04 | 28.68 | 27.38 | 30.04 | **<2.2E-308** |  |  |  |  |
| DCD | Males | 1.86 | 17.90 | 16.38 | 19.57 | **<2.2E-308** | 0.97 | 0.82 | 1.14 | 0.71 |
|  | Females | 1.08 | 17.61 | 15.25 | 20.33 | **<2.2E-308** |  |  |  |  |
| ID | Males | 5.30 | 9.82 | 9.38 | 10.28 | **<2.2E-308** | 1.11 | 1.03 | 1.20 | 0.0090 |
|  | Females | 4.89 | 10.89 | 10.23 | 11.60 | **<2.2E-308** |  |  |  |  |
| Epilepsy | Males | 2.57 | 3.27 | 3.09 | 3.47 | **<2.2E-308** | 1.08 | 0.99 | 1.19 | 0.099 |
|  | Females | 2.85 | 3.55 | 3.29 | 3.82 | **6.6E-238** |  |  |  |  |
| Congenital malformations | Males | 8.20 | 1.40 | 1.36 | 1.45 | **1.8E-109** | 1.11 | 1.05 | 1.17 | **2.5E-04** |
|  | Females | 6.46 | 1.54 | 1.47 | 1.62 | **8.9E-72** |  |  |  |  |
| Chromosomal abnormalities | Males | 0.60 | 2.96 | 2.63 | 3.34 | **2.7E-70** | 0.84 | 0.68 | 1.03 | 0.096 |
|  | Females | 0.52 | 2.48 | 2.07 | 2.96 | **9.5E-24** |  |  |  |  |

Abbreviations: ADHD: attention-deficit/hyperactivity disorder; ASD: autism spectrum disorder; DCD: developmental coordination disorder; ID: intellectual disability. Sex is coded as 0 for male and 1 for female. Birth year is included as a covariate. Estimated p-values below a Bonferroni-adjusted threshold for multiple testing ( $p < 0.0083$ ) are bolded.

### Testing the female protective effect hypothesis

#### Polygenic risk score analysis

Results of meta-analyses of each leave-one-study out logistic regression analysis for ADHD PRS are shown in Figure 4. There was no association of ADHD PRS with sex in cases (OR=1.02 [95% CI: 0.98-1.06], p=0.28, mean R^2^ [SE]=0.0019 [0.00039]). Sensitivity tests were run excluding the data from 23andMe and all non-European ancestry individuals from the discovery sample and then additionally not including sex as a covariate in the discovery GWAS analyses; results remained similar (Figure S7). There was also no association of ADHD PRS with sex in controls (OR=0.99 [95% CI: 0.96-1.01], p=0.23, mean R^2^ [SE]=0.0011 [0.00024]; Figure S8).

**Figure 4.**
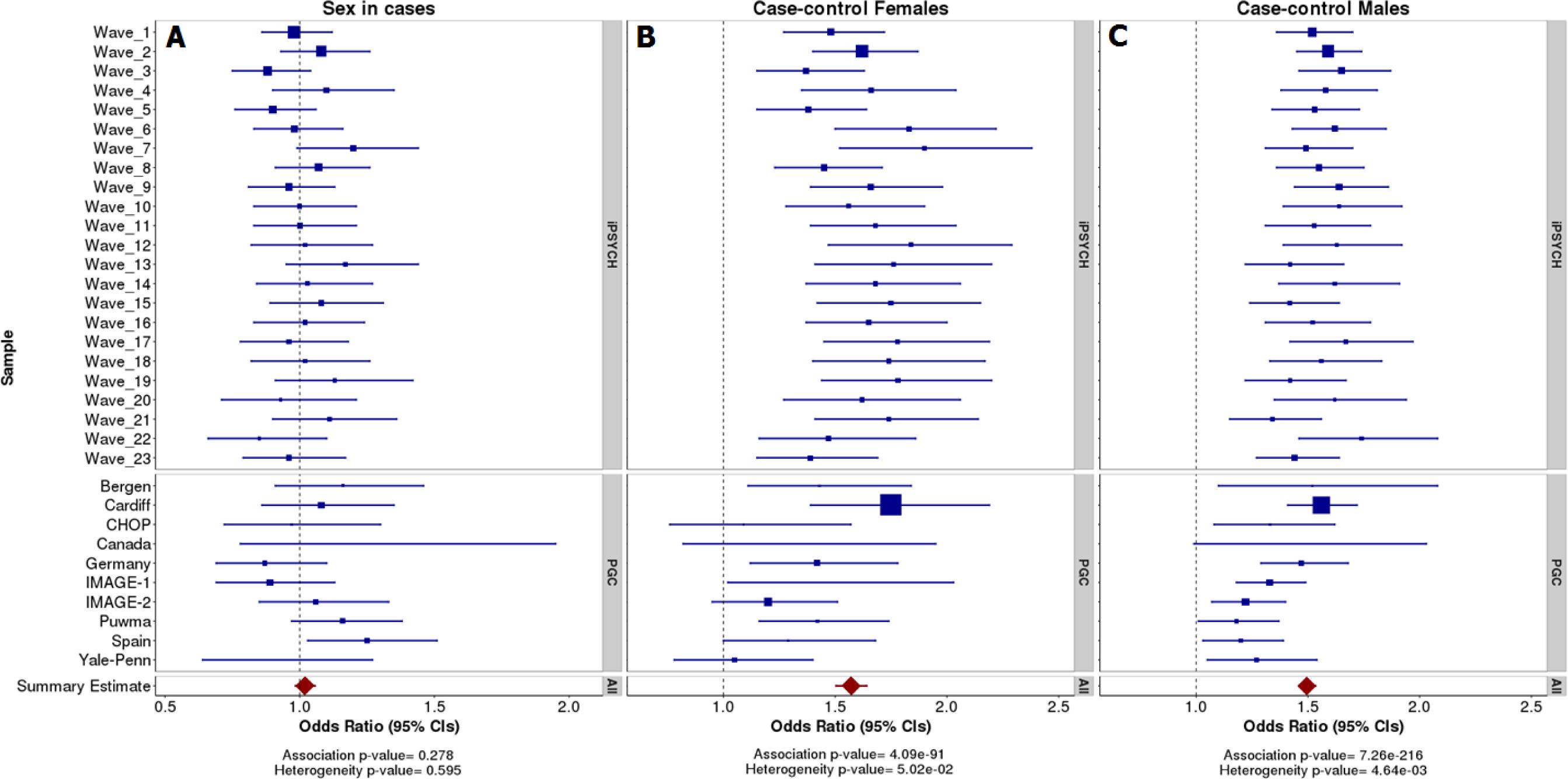
Forest plots of meta-analysis results for logistic regression analyses of ADHD polygenic risk score with: a) case sex as the outcome; b) case-control status in females; c) case-control status in males.

There was a clear association of ADHD PRS with ADHD case status in both sex-specific analyses. Similar differences in PRS between cases and controls were observed for females (OR=1.57 [95% CI: 1.50-1.64], p=4.1xE-91, mean R^2^ [SE]=0.039 [0.0034]) and males (OR=1.50 [95% CI: 1.46-1.53], p=7.3xE-216, mean R^2^ [SE]=0.032 [0.0024]). Several of the PGC studies did not show a significant association with case-control status in just females, but given that these samples had relatively few females (with the lowest N of 27 female case/pseudo-control pairs in the Canadian sample; see Online Methods) this is likely to be due to very low power in these studies.

#### Epidemiological analysis

To test for evidence of increased risk for ADHD in siblings of females with ADHD, we used Swedish population data to select sibling pairs where at least one child had ADHD. We stratified the comparison siblings by sex and tested whether the siblings of female probands were at increased risk of ADHD as compared to siblings of male probands. The results showed that having a female sibling diagnosed with ADHD is associated with an increased risk for an ADHD diagnosis in males (OR=1.18, [95% CI: 1.13-1.23], p=1.8E-13) and females (OR=1.09, [95% CI: 1.01-1.16], p=0.017). Co-varying for presence of ASD, congenital malformations, and ID in the proband did not affect the results (OR_MALE_=1.18, [95% CI: 1.13-1.23], p=1.4E-13; OR_FEMALE_=1.09, [95% CI: 1.02-1.16], p=0.014).

## Discussion

We tested two specific hypotheses for the male bias in ADHD: first, that sex-specific genetic heterogeneity may be linked with clinical heterogeneity, affecting prevalence rates, and second, that females affected with ADHD may be carriers of an increased burden of genetic risk variants, as compared with affected males. We analyzed common variant autosomal data from the largest available ADHD case-control GWAS sample (N=55,374 individuals) and Swedish population cohort data from nearly 2 million individuals. Using genome-wide analyses of autosomal common variants, we demonstrated a high level of genetic correlation for ADHD in males and females and found no clear increase of polygenic burden in affected females compared to affected males. However, we did observe, in a different sample, that siblings of females with ADHD are at an increased risk of having ADHD, compared to siblings of affected males. The results from the epidemiological sample also suggested that females diagnosed with ADHD may be at an especially high risk of certain comorbid developmental conditions compared to affected males.

The observed high SNP-based genetic correlation in the tests of sex-specific heterogeneity suggest that, to a large extent, the same common autosomal genetic risk variants are involved in ADHD for both sexes. The results are consistent across 2 methods, although the analysis of PGC-only data using LDSC is somewhat lower, possibly because LDSC is less well optimized for analyses of smaller samples. Our conclusion is strengthened by the observations that including sex as a covariate had a minimal effect on the overall GWAS results for ADHD and that no loci showed significant heterogeneity across the sexes or a sex-by-genotype interaction. While sex-specific genetic heterogeneity from common autosomal variants seems unlikely based on our results, this study cannot rule out the possibility that heterogeneous effects could exist for rare or non-autosomal variation or that with increased sample sizes, weaker effects of common variant genetic heterogeneity could be detected.

Indeed, the epidemiological analyses of Swedish population data do suggest that some degree of clinical and/or etiological heterogeneity does exist. ADHD was associated with comorbid diagnoses of ASD, DCD, ID, epilepsy, congenital malformations, and chromosomal abnormalities in both males and females. ADHD-by-sex interaction analyses revealed that the strength of association was greater in females for ASD, congenital malformations, and (to a lesser extent) also for mild ID. There may be several possible explanations for these findings. First, females with ADHD may have a higher than expected risk of comorbid severe conditions and as such, may have a higher level of clinical heterogeneity, as compared to males with ADHD. Alternatively, ascertainment and diagnostic biases, where females are more likely to be diagnosed with ADHD if they have a more severe phenotypic presentation, could be involved.

In the first case, these results could indirectly point to a greater role of rare, deleterious genetic variants in females with ADHD, as such rare variants are strongly implicated in some of the comorbid conditions assessed^12–22^. Furthermore, there is evidence for such an imbalance in other neurodevelopmental disorders, as females with ASD and developmental delay show a consistently increased burden of disruptive CNVs and rare, deleterious single nucleotide mutations compared with affected males^12–16^. However, this line of thinking needs to be tested directly, as common genetic variants also play an important role in complex disorders, such as ASD^32,33^, and too little is currently known about the contribution of rare genetic variants to ADHD, stratified by sex.

Although an increased burden of rare variants in ADHD females is possible, our results are inconclusive. When examining rare genetic syndromes linked to chromosomal abnormalities (including autosomal variants and sex chromosome aneuploidies), we see no evidence of an ADHD-by-sex interaction. It is also unclear why the results show a stronger association in females between ADHD and mild ID but not for more severe degrees of ID. If females diagnosed with ADHD are indeed enriched for rare deleterious mutations, one interesting secondary question is what the clinical presentation of males with rare deleterious variants is; do they present with a more complex phenotype, without a diagnosis of ADHD, or are such individuals less likely to survive past early childhood? Whilst there is some evidence that there are overall sex differences in the population, with females generally more likely to harbor large, rare CNVs^34^, it is too early to draw definitive conclusions.

Furthermore, a limitation of the epidemiological analyses is the possibility of ascertainment and diagnostic biases. Individuals who receive one diagnosis become the focus of clinical attention and are more likely to receive subsequent diagnoses, whereas individuals with less complex phenotypes might not come to clinical attention. If females are routinely under-diagnosed with ADHD or other neurodevelopmental disorders, this issue may disproportionately affect ascertainment of female cases, leading to the observed pattern of results. Other possible sources of bias include typical exclusion criteria for diagnosing ADHD (e.g. ID or ASD) and the possibility of an inflated false positive rate of diagnoses due to diagnostic uncertainty and change over time. We endeavored to limit the impact of the latter issue by only considering diagnoses in individuals who had at least 2 reported diagnoses for any of the studied conditions. Although the pattern of results is consistent with the possibility of increased clinical and etiological heterogeneity in females with ADHD, genetic studies of rare variation are required to rule out these alternative explanations for our findings.

Epidemiological analyses also showed that siblings of females diagnosed with ADHD were at higher risk of being diagnosed with ADHD than siblings of diagnosed males. This confirms results from previous family studies^26–28^, supporting the hypothesis that females require a greater burden of genetic risk to manifest ADHD. However, the effect sizes were not large (OR=1.09 – 1.18), suggesting that an increased burden of inherited genetic variation may only be a small contribution to the sex bias in ADHD prevalence. These results could also occur if clinicians had a higher threshold for diagnosing ADHD in females or were more likely to diagnose it in females if accompanied by a comorbid disorder, although we did not see an attenuation of our results when comorbid conditions in the proband were accounted for in analyses.

Contrary to previous smaller studies^3,31^, we did not find an association between ADHD PRS and case sex; no enrichment of polygenic burden from common variants was observed in females with ADHD. Analyses in ASD are consistent with the current study by not finding an increased burden from common variants in affected females^35,36^, in contrast to rare variant studies (see above and^12–16^).

One possible explanation for the observed results is that a higher degree of genetic heterogeneity within females may have masked any differences in PRS burden by sex in the current study. It has been suggested that common and rare variants contribute additively to risk of ADHD, with cases who are non-CNV carriers having lower ADHD PRS than cases who have large, rare CNVs^37^. Thus, if females diagnosed with ADHD are more likely to have a more complex syndromic phenotype (as suggested by the Swedish population analyses) and given that such phenotypes are more likely to be associated with rare variants^12–22^, this subgroup of females could have on average lower PRS than males with ADHD. On the other hand, affected females with a less severe phenotypic presentation, who are not carriers of such rare variants, could have higher PRS than affected males. If this were the case, any overall difference in PRS between the sexes could be obscured. Rare variant data are needed together with common variant data to address this possibility.

Although the focus of this manuscript was on possible genetic sources of influence on the sex bias in ADHD, other factors, such as ascertainment and diagnostic biases, may play an important role. There is some indication that females are more likely to be diagnosed with the predominantly inattentive subtype of ADHD and present with inattentive symptoms, whereas males are more likely to be diagnosed with the combined subtype of ADHD and present with hyperactive-impulsive symptoms and comorbid disruptive behavioral problems^38–41^. Relative prevalence rates also vary by diagnostic instrument used and case ascertainment. For example, the ratio of male to female cases in the Swedish population was 2:1, somewhat lower than in the iPSYCH Danish population (2.8:1) and PGC clinical (3.5:1) samples. ADHD cases with diagnosed moderate-severe ID (IQ<50) were excluded from iPSYCH. ADHD cases in the PGC studies were primarily ascertained from clinics, ADHD was confirmed with structured interviews and children with comorbid ASD, epilepsy, ID (IQ<70) and other conditions were excluded. As such, the false positive rate for an ADHD diagnosis is likely to be higher in the iPSYCH and Swedish registry-based datasets than in the PGC, while the latter is likely under-represented for individuals with ADHD neurodevelopmental co-morbidities. Another major difference is that many of the PGC studies utilized DSM criteria and thus included children with inattentive and hyperactive-impulsive subtypes of ADHD, whereas the Swedish, iPSYCH (and some of the European PGC studies) used the stricter ICD definition. Despite these differences, we find very high genetic correlation between PGC and iPSYCH, suggesting that overall these diagnostic differences do not have a perceptible impact on the involvement of common risk variants. We saw a similar pattern of results for PGC-only and iPSYCH-only in the sex-specific analyses, with the caveat that the PGC study was somewhat smaller and thus confidence intervals were larger.

Another possible contribution to sex bias that was beyond the scope of the current study is the role of sex hormones and sex chromosomes. There is broad evidence for a specific role of sex hormones (e.g. estrogen) and sex chromosomes (e.g. X chromosome aneuploidy) on early brain development and neurodevelopmental disorders such as ADHD^25,42,43,^ suggesting that future efforts to comprehensively examine the role of the sex chromosomes and their downstream products in the male bias in ADHD may be worthwhile.

The results of this study demonstrate a clear polygenic contribution from common autosomal genetic variants to ADHD in both sexes, as evidenced by the moderate SNP-h^2^ estimates using two methods, the clear deviation of test statistics on sex-specific GWAS QQ-plots, and the significantly higher overall PRS in cases of either sex when compared with sex-matched controls. Top hits from sex-specific GWAS analyses corroborate the results from a combined analysis of both sexes^7^.

The results of the high genetic correlation between male and female cases with ADHD support combining GWAS data from both sexes in meta-analyses of ADHD and further suggest that current clinical practices of diagnosing ADHD are capturing a clinical phenotype that is similar at the level of common genetic risk variants in both sexes. The results of epidemiological analyses do suggest some degree of clinical heterogeneity, with ADHD showing a stronger association with comorbid ASD, congenital malformations, and possibly also mild ID in females. Although we find evidence for an overall increased burden of inherited genetic risk variants for ADHD in females based on sibling analysis, there is no difference in polygenic burden in males and females, as measured by PRS. Further work simultaneously examining the role of variants across the spectrum of frequencies is needed to comprehensively examine the role of genetic risk in the sex bias in ADHD prevalence.

## Online Methods

### Description of Genetic data

Genotype data for ADHD cases and control individuals were available from the Psychiatric Genomics Consortium (PGC) and the Lundbeck Foundation Initiative for Integrative Psychiatric Research (iPSYCH). See GWAS publication for full details^7^. The PGC ADHD samples came from a range of studies that were predominantly of European ancestry. They consisted of clinically-ascertained cases of ADHD matched with either controls from the same ancestry group or with pseudo-controls created from the non-transmitted alleles of both parents (trio samples). The individual studies have been previously described in more detail in individual publications^6,44–57^. The iPSYCH sample is based on genotyping of Guthrie cards obtained from the Danish Neonatal Screening Biobank. Blood-spot samples were collected and frozen shortly after birth for individuals born in Denmark and stored in the Danish Newborn Screening Biobank and Statens Serum Institute. The individuals included in the iPSYCH sample were born between May 1, 1981 and December 31, 2005, and had to be alive and resident in Denmark after one year and have a known mother. Cases with ADHD diagnoses (ICD-10 code F90.0) were identified using the Danish Psychiatric Central Research Register. This register includes data on everyone admitted to a psychiatric hospital for assessment or treatment (between 1969 and 2013), as well as everyone who attended psychiatric outpatient services (between 1995 and 2013). Control individuals were randomly selected from the population. The DNA from these samples was extracted, whole-genome amplified in triplicates and genotyped in 23 batches (referred to from here on as waves) using the Illumina PsychChip (a customized HumanCoreExome chip). The first wave consists of the youngest samples (born in 2004) and wave 23 consists of the oldest samples (born in 1981). The study was approved by the Danish Data Protection Agency and the Scientific Ethics Committee in Denmark.

Summary statistics from a GWAS of self-reported ADHD including sex as a covariate were also available from the personal genetics company 23andMe, Inc. Research participants of 23andMe provided informed consent and participated in research online, under a protocol approved by the external AAHRPP-accredited IRB, Ethical & Independent Review Services (E&I Review). The GWAS was based on data from 5,857 self-assessed ADHD cases and 70,393 controls and had a genetic correlation of 0.653 (0.114) with the PGC+iPSYCH samples^7^. Results from this GWAS were only used for the polygenic risk score analyses as no raw genotypes or sex-specific summary data were available.

### Quality control and data preparation

PGC and iPSYCH samples were processed using the Rapid Imputation Consortium Pipeline (Ricopili), which is a quality control (QC), imputation, and principal components analysis (PCA) pipeline developed and used by the PGC and collaborators. See GWAS publication for full details^7^. QC, imputation and PCA were performed separately using the PGC pipeline (Ricopili) for each PGC study and the 23 waves of iPSYCH samples, with the exception that PCA was performed in the entire iPSYCH sample simultaneously. The 1000 Genomes Project, phase 3, data were used as the imputation reference. Cross-study (PGC) and cross-wave (iPSYCH) relatedness analyses were performed in PLINK-v.1.9 on merged, LD-pruned datasets. One of each pair of individuals related at Π > 0.2 was excluded (preferentially keeping cases over controls).

See Table S8 for sex-stratified sample sizes for each PGC study and iPSYCH wave. The total sample size after all quality control was N=20,183 cases (25% females) and N=35,191 pseudo-controls/population controls (38% females). Analyses that were restricted to European-only samples consisted of 19,099 cases (26% females) and 34,194 controls (38% females). ADHD GWAS summary statistics were also available from research participants of the personal genetics company 23andMe, Inc. (N=5,857 self-reported ADHD cases, 70,393 controls).

### Sex-specific GWAS analyses

Sex-specific case-control genome-wide logistic regression analyses of imputed autosomal dosage data were performed in each PGC study and iPSYCH wave separately, using the “--dosage” option in PLINK-version-1.9, co-varying for principal components (PCs) and/or site indicator variables, as appropriate. iPSYCH samples included the first 4 PCs and any PCs significantly associated with case status, obtained from the joint PCA in the entire iPSYCH sample. For PGC studies with <1000 samples, the top 5 PCs were used and for studies with ≥1000 samples, the first 10 PCs were used as covariates. For the IMAGE-1 study, indicator variables coding for site ID were included as covariates instead of PCs, as this study used a trio design but consisted of samples contributed by several different data collection sites. Trio studies were split by case sex, keeping each pseudo-control together with its corresponding case.

Results were filtered for each study/wave and SNPs meeting the following criteria were retained for the sex-specific analyses: imputation quality (INFO score) > 0.8, call rate in best guess genotype data > 0.925, minor allele frequency (MAF) > 0.01, and expected MAF in cases (2 x MAF in controls x no. of cases) > 1. Sex-specific GWAS meta-analyses of filtered results were performed in METAL using the standard error analysis scheme (STDERR). Meta-analysis results were additionally filtered to retain only SNPs that were available for analysis in at least half of the total sample size and present in both the male-only and female-only analyses. This yielded results for N=7,531,543 common variants in the meta-analyses (hereafter: PGC+iPSYCH).

### Estimating SNP-heritability and genetic correlation

Bivariate LD score regression (using LDSC^58,59^) analyses were run on the sex-specific meta-analyzed summary statistics. The primary analyses (with the most power) are those for the full PGC+iPSYCH sample but we also examined estimates in the PGC and iPSYCH samples separately using LDSC and a second method, GREML (using GCTA^60,61^), to examine the stability of the findings. Sex-specific heritability was also estimated using univariate models. Analyses were restricted to European-only samples.

LD scores from a European reference panel provided with the LDSC software were used for analysis. LDSC analyses were based on the following numbers of SNPs, after restriction to HapMap SNPs: PGC-only: 1,108,369 SNPs; iPSYCH-only: 1,021,086 SNPs; PGC+iPSYCH 1,023,856 SNPs. The intercept was not constrained in LDSC, to provide unbiased estimation. For sensitivity, genetic correlation analyses were also run in LDSC to assess cross-dataset (PGC vs. iPSYCH) within- and across-sex genetic correlations.

Because of strict restrictions on access to individual genotypes, bivariate GREML analyses were only performed separately in the PGC and iPSYCH samples. For each of these datasets, best guess genotype data were generated using Ricopili and strictly filtered (MAF>0.05, in addition to previous frequency, imputation quality and other filters). Genotypes were merged together across studies using PLINK. Asymmetric/ambiguous (AT, TA, CG, GC), multi-allelic and duplicate position SNPs were excluded. For each dataset, a genomic-relationship matrix was calculated using GCTA, restricted to HapMap-3 SNPs. Analyses were based on the following numbers of SNPs: PGC-only: 191,466 SNPs; iPSYCH-only: 435,086 SNPs. One of each pair of individuals related at the level of second cousins (pi-hat>0.05) was excluded, preferentially keeping cases; this excluded: N=16 cases and N=91 controls in the PGC dataset and N=^1^,439 cases and N=3,170 controls in the iPSYCH dataset. PCA (after LD-pruning and removing SNPs located in long-range LD regions) was performed on the merged, unrelated samples using PLINK, to derive population covariates. The first 10 PCs as well as binary study/wave indicators were used as covariates for subsequent analyses. Bivariate GREML was used to estimate the genetic correlation across-sex. Univariate GREML analyses in GCTA were used to estimate SNP-h^2^ in males and females with ADHD relative to sex-matched controls.

The expected range of the genetic correlation (r_g_) estimates should be from -1 to 1. However, the estimator was left unconstrained for these analyses in GREML and LDSC to allow for an unbiased assessment of the standard errors of the estimates; as such, some of the estimates exceed this range. Specific tests were used to determine whether the SNP-h^2^ (on the liability scale) estimates differed significantly for males and females using the formula: (SNP-h^2^_F_ – SNP-h^2^_M_)^2^/(SE_F_^2^ + SE_M_^2^) with a Chi^2^ test with 1 degree of freedom. One-tailed tests were also used to determine whether the estimates of genetic correlation differed significantly from one (z=(1-r_g_)/SE) or from zero (z=r_g_)/SE), compared to a normal distribution.

Based on an estimated population prevalence rate of approximately 5%^1^ for ADHD and an observed male:female ratio of approximately 3:1 in the cases, the following prevalence rates were assumed for converting the estimates of SNP-heritability to the liability scale: 2.5% in females and 7.5% in males. Analyses were also re-run assuming different relative population prevalence for males and females, depending on the assumed ratio of the relative prevalence (ranging from equal prevalence assumed to a 7:1 male bias). This was done to examine the sensitivity of this assumption on the estimation of liability scale SNP-h^2^. These analyses were also repeated while randomly down-sampling the number of male cases and controls to match the available sample size for females within each study/wave.

### Secondary GWAS analyses

A number of secondary GWAS analyses were run to further examine the impact of sex on genome-wide association analyses of ADHD. First, heterogeneity statistics from a meta-analysis of the male-only and female-only summary statistics were examined for all SNPs. Second, combined GWAS analyses including a sex-by-genotype interaction term were carried out. Third, the genetic correlation was estimated using LDSC for GWAS analyses of the combined sample including and excluding sex as a covariate. Finally, GWAS analyses of case sex (male cases coded as 0 and female cases coded as 1) were carried out.

### Polygenic risk score analyses

A leave-one-study/wave-out approach was used to maximize power and maintain fully independent target and discovery samples for polygenic risk score (PRS) analyses, using the standard approach^62,63^. First, GWAS analyses of imputed dosage data were run for all samples in each PGC study and iPSYCH wave separately, as described previously, co-varying for PCs as well as sex. Meta-analyses using METAL (with the STDERR scheme) were run excluding one set of summary results at a time, for each combination of studies. To maximize power for the discovery samples, GWAS results from 23andMe and non-European samples were also included in the ADHD discovery meta-analyses. For each set of discovery results, LD-clumping was run to obtain a relatively independent set of SNPs, while retaining the most significant SNP in each LD block. The following parameters were applied in PLINK: --clump-kb 500 -- clump-r2 0.3 --clump-p1 0.5 --clump-p2 0.5. Asymmetric/ambiguous (AT, TA, CG, GC) SNPs, indels, multi-allelic and duplicate position SNPs were excluded. The SNP selection p-value threshold used was p<0.1. The number of clumped SNPs for each study/wave varied from 20596-43427 (see Table S9).

PRS were calculated for each individual in the independent target sample (restricted to European samples) by scoring the number of risk alleles (weighted by the SNP log of the odds ratio) across the set of clumped, meta-analyzed SNPs in PLINK.v.1.9 (using the command -- score no-mean-imputation). Scores were derived in best guess genotype data after filtering out SNPs with MAF<0.05 and INFO<0.8. The PRS were standardized using z-score transformations; odds ratios can be interpreted as the increase in risk of the outcome, per standard deviation in PRS. Logistic regression analyses including PCs tested for association of PRS with sex in the cases (males were coded as 0 and females were coded as 1) and case status, stratified by sex. Finally, overall meta-analyses of the leave-one-out analyses were performed. The analyses were re-run using European-only samples and then also by excluding sex as a covariate in the discovery GWAS analyses, as sensitivity tests. All regression and meta-analyses were run in R-3.2.2.

### Epidemiological analyses

Analyses of Swedish registry data were based on all individuals born in Sweden between 1987 and 2006, as identified using the Medical Birth Register. Data linkage of several nation-wide Swedish registers was performed using the unique personal identification number^64^. Information from the Total Population Register^65^, Cause of Death Register and the Multi-Generation Register^66^ were used to identify those individuals of known maternity and paternity who lived in Sweden at least until age 12 years (or until the time of this study, if they were younger than 12 years old). Information on ADHD diagnoses was obtained from the National Patient Register^67^ for ICD-9 (1987-1996) and ICD-10 (1997-2013) and from the Prescribed Drug Register^68^ (June 2005-2014). ADHD cases were defined as those individuals who had at least 2 recorded diagnoses of ADHD or 2 recorded prescriptions of ADHD medication (Methylphenidate, Amphetamine, Dexamphetamine, Atomoxetine or Lisdexamfetamine) after the age of 3 years. Analyses were based on N=77,905 ADHD cases and N=1,874,637 control individuals. The data linkage of the Swedish registry data was approved by the regional ethics review board in Stockholm, Sweden. The requirement for informed consent was waived, because the study was register-based, and individuals were not personally identifiable at any time.

All epidemiological analyses were performed in R (with the ‘drgee’ package). Children were clustered together if they shared the same biological mother, in order to obtain standard errors that accounted for non-independent observations. Birth year was included as a covariate in all analyses.

We assessed whether females affected with ADHD are at a higher risk than males for comorbid severe developmental disorders and rare genetic syndromes. International Classification of Diseases (ICD) codes for the following categories of disorders were examined: intellectual disability (ID), autism spectrum disorder (ASD), developmental coordination disorder (DCD), epilepsy, congenital malformations (CM) and chromosomal abnormalities (CA); see Table S10 for specific ICD codes. Diagnoses of ASD and DCD were only considered after age 1 year and ID diagnoses after age 2 years. No age restrictions were made for epilepsy, CM or CA. For each comorbid condition, individuals were considered as affected (coded as 1) if they had at least 2 recorded diagnoses in that category and unaffected (coded as 0) if they did not meet these criteria. Generalized estimating equations were used to test for the effect of an ADHD-by-sex interaction term on each outcome. First, the effect of presence of ADHD on each outcome within individuals was estimated separately for males and females (analytic model: gee(outcome ~ ADHD + birth_year)). Next, we tested for an ADHD-by-sex interaction term on each outcome, using the following analytic model: gee(outcome ~ ADHD + sex + ADHD*sex + birth_year). For individuals with available information on severity of ID, secondary analyses were run for 3 severity categories: mild, moderate and severe/profound.

To test the female protective effect hypothesis, we estimated whether risk of ADHD in siblings of females with ADHD was higher than for siblings of affected males, stratified by the sex of the comparison sibling. Analyses were restricted to pairs of full siblings, based on sharing both biological parents). Twins (i.e. children born with 2-weeks of each other) were excluded as zygosity could not be confirmed. Analyses were restricted to sibling pairs with at least 1 child who had a diagnosis of ADHD, as defined above (N=71,691 observations (of which, N=23,452 came from female probands), consisting of N=21,784 unique index individuals, of which N=7,186 came from unique female probands). The effect of the proband being female on the comparison sibling’s risk for ADHD was estimated using the following model, stratified by the sex of the comparison sibling: gee(ADHD_sib2 ~ sex_sib1 + birth_year_sib2).

## Acknowledgements & Funding

Dr J. Martin was supported by the Wellcome Trust (Grant No: 106047).

The Broad Institute and Massachusetts General Hospital investigators would like to acknowledge support from the Stanley Medical Research Institute and NIH grants: 1R01MH094469 (PI: Neale), 1R01MH107649-01 (PI: Neale).

The iPSYCH team acknowledges funding from the Lundbeck Foundation (grant no R102-A9118 and R155-2014-1724), the Stanley Medical Research Institute, the European Research Council (project no: 294838), the European Community’s Horizon 2020 Programme (H2020/2014-2020) under Grant No. 667302 (CoCA), the Novo Nordisk Foundation for supporting the Danish National Biobank resource, and grants from Aarhus and Copenhagen Universities and University Hospitals, including support to the iSEQ Center, the GenomeDK HPC facility, and the CIRRAU Center.

We would like to thank the research participants and employees of 23andMe for making this work possible. This work was supported by the National Human Genome Research Institute of the National Institutes of Health (grant number R44HG006981).

## Consortium Members

Please see the Supplementary Note for a list of members and affiliations

## Author contributions

JM contributed to study conception, data analysis, interpretation of the results and drafting the manuscript. RW, DD, MM contributed to data analysis, interpretation of the results and critical revision of the manuscript. AT, MO, BN contributed to study conception, interpretation of the results and critical revision of the manuscript. IB, LG contributed to data analysis and critical revision of the manuscript. SHL, NW contributed to interpretation of the results and critical revision of the manuscript. ADB contributed to data generation and design of the overall project and critical revision of the manuscript. TW, PBM, MGP, OM, MN, DMH, JBG contributed to data generation and design of the overall project. HL, PL, ER, NE, BF, SF contributed to critical revision of the manuscript.

## References

1. Polanczyk, G., de Lima, M. S., Horta, B. L., Biederman, J. & Rohde, L. A. The worldwide prevalence of ADHD: a systematic review and metaregression analysis. Am. J. Psychiatry 164, 942 (2007).

2. Faraone, S. V. et al. Attention-deficit/hyperactivity disorder. Nat. Rev. Dis. Prim. 1, 15020 (2015).

3. Hamshere, M. L. et al. High loading of polygenic risk for ADHD in children with comorbid aggression. Am. J. Psychiatry 170, 909–916 (2013).

4. Cross-Disorder Group of the Psychiatric Genomics Consortium. Genetic relationship between five psychiatric disorders estimated from genome-wide SNPs. Nat. Genet. 45, 984–94 (2013).

5. Williams, N. M. et al. Genome-wide analysis of copy number variants in attention deficit hyperactivity disorder: the role of rare variants and duplications at 15q13. 3. Am. J. Psychiatry 169, 195–204 (2012).

6. Yang, L. et al. Polygenic transmission and complex neuro developmental network for attention deficit hyperactivity disorder: Genome-wide association study of both common and rare variants. Am. J. Med. Genet. Part B Neuropsychiatr. Genet. 162B, 419–430 (2013).

7. Demontis, D. et al. Discovery Of The First Genome-Wide Significant Risk Loci For ADHD. bioRxiv (2017). doi: https://doi.org/10.1101/145581

8. Lahey, B. B., Applegate, B., McBurnett, K. & Biederman, J. DMS-IV field trials for attention deficit hyperactivity disorder in children and adolescents. Am. J. Psychiatry 151, 1673–1685 (1994).

9. Polderman, T. J. C. et al. Meta-analysis of the heritability of human traits based on fifty years of twin studies. Nat. Genet. 47, 702–709 (2015).

10. Weiss, L. A., Pan, L., Abney, M. & Ober, C. The sex-specific genetic architecture of quantitative traits in humans. Nat. Genet. 38, 218–22 (2006).

11. Yang, J. et al. Genome-wide genetic homogeneity between sexes and populations for human height and body mass index. Hum. Mol. Genet. 24, 7445–9 (2015).

12. Gilman, S. R. et al. Rare de novo variants associated with autism implicate a large functional network of genes involved in formation and function of synapses. Neuron 70, 898–907 (2011).

13. Levy, D. et al. Rare de novo and transmitted copy-number variation in autistic spectrum disorders. Neuron 70, 886–897 (2011).

14. Neale, B. M. et al. Patterns and rates of exonic de novo mutations in autism spectrum disorders. Nature 485, 242–245 (2012).

15. Iossifov, I. et al. De novo gene disruptions in children on the autistic spectrum. Neuron 74, 285–299 (2012).

16. Jacquemont, S. et al. A Higher Mutational Burden in Females Supports a ‘Female Protective Model’ in Neurodevelopmental Disorders. Am. J. Hum. Genet. 94, 415–425 (2014).

17. Girirajan, S. et al. Phenotypic heterogeneity of genomic disorders and rare copy-number variants. N. Engl. J. Med. 367, 1321–1131 (2012).

18. Pescosolido, M. & Gamsiz, E. Distribution of disease-associated copy number variants across distinct disorders of cognitive development. J.… 52, 414–430.e14 (2013).

19. Guilmatre, A. et al. Recurrent rearrangements in synaptic and neurodevelopmental genes and shared biologic pathways in schizophrenia, autism, and mental retardation. Arch. Gen. Psychiatry 66, 947 (2009).

20. Iossifov, I. et al. The contribution of de novo coding mutations to autism spectrum disorder. Nature 515, 216–221 (2014).

21. De Rubeis, S. et al. Synaptic, transcriptional and chromatin genes disrupted in autism. Nature 515, 209–215 (2014).

22. Samocha, K. E. et al. A framework for the interpretation of de novo mutation in human disease. Nat. Genet. 46, 944–950 (2014).

23. Williams, N. M. et al. Rare chromosomal deletions and duplications in attention-deficit hyperactivity disorder: a genome-wide analysis. Lancet 376, 1401–1408 (2010).

24. Vorstman, J. A. S. S. & Ophoff, R. A. Genetic causes of developmental disorders. Curr. Opin. Neurol. 26, 128–36 (2013).

25. Scerif, G. & Baker, K. Annual Research Review: Rare genotypes and childhood psychopathology - uncovering diverse developmental mechanisms of ADHD risk. J. Child Psychol. Psychiatry 56, 251–273 (2015).

26. Rhee, S. H. & Waldman, I. D. Etiology of sex differences in the prevalence of ADHD: An examination of inattention and hyperactivity–impulsivity. Am. J. Med. Genet. Part B Neuropsychiatr. Genet. 127, 60–64 (2004).

27. Taylor, M. J. et al. Is There a Female Protective Effect Against Attention-Deficit/Hyperactivity Disorder? Evidence From Two Representative Twin Samples. J. Am. Acad. Child Adolesc. Psychiatry (2016). doi: 10.1016/j.jaac.2016.04.004

28. Smalley, S. L. et al. Familial clustering of symptoms and disruptive behaviors in multiplex families with attention-deficit/hyperactivity disorder. J. Am. Acad. Child Adolesc. Psychiatry 39, 1135–1143 (2000).

29. Faraone, S. V. Family Study of Girls With Attention Deficit Hyperactivity Disorder. Am. J. Psychiatry 157, 1077–1083 (2000).

30. Chen, Q. et al. Familial aggregation of attention-deficit/hyperactivity disorder. J. Child Psychol. Psychiatry (2016). doi: 10.1111/jcpp.12616

31. Martin, J. et al. Genetic risk for attention-deficit/hyperactivity disorder contributes to neurodevelopmental traits in the general population. Biol. Psychiatry 76, 664–71 (2014).

32. Gaugler, T. et al. Most genetic risk for autism resides with common variation. Nat. Genet. 46, 881–885 (2014).

33. Klei, L. et al. Common genetic variants, acting additively, are a major source of risk for autism. Mol. Autism 3, 9 (2012).

34. Han, J. et al. Gender differences in CNV burden do not confound schizophrenia CNV associations. Sci. Rep. 6, 25986 (2016).

35. Mitra, I. et al. Pleiotropic Mechanisms Indicated for Sex Differences in Autism. PLOS Genet. 12, e1006425 (2016).

36. Weiner, D. J. et al. Polygenic transmission disequilibrium confirms that common and rare variation act additively to create risk for autism spectrum disorders. Nat. Genet. (2017). doi: 10.1038/ng.3863

37. Martin, J., O’Donovan, M. C., Thapar, A., Langley, K. & Williams, N. The relative contribution of common and rare genetic variants to ADHD. Transl. Psychiatry 5, e506 (2015).

38. Biederman, J. et al. Influence of Gender on Attention Deficit Hyperactivity Disorder in Children Referred to a Psychiatric Clinic. Am. J. Psychiatry 159, 36–42 (2002).

39. Weiss, M., Worling, D. & Wasdell, M. A chart review study of the inattentive and combined types of ADHD. J. Atten. Disord. 7, 1–9 (2003).

40. Willcutt, E. G. The prevalence of DSM-IV attention-deficit/hyperactivity disorder: a meta-analytic review. Neurother. J. Am. Soc. Exp. Neurother. 9, 490–9 (2012).

41. Staller, J. & Faraone, S. V. Attention-Deficit Hyperactivity Disorder in Girls. CNS Drugs 20, 107–123 (2006).

42. Printzlau, F., Wolstencroft, J. & Skuse, D. H. Cognitive, behavioral, and neural consequences of sex chromosome aneuploidy. J. Neurosci. Res. 95, 311–319 (2017).

43. Loke, H., Harley, V. & Lee, J. Biological factors underlying sex differences in neurological disorders. Int. J. Biochem. Cell Biol. 65, 139–150 (2015).

44. Neale, B. M. et al. Genome-wide association scan of attention deficit hyperactivity disorder. Am. J. Med. Genet. (Neuropsychiatric Genet. 147B, 1337–1344 (2008).

45. Neale, B. M. et al. Meta-analysis of genome-wide association studies of attention-deficit/hyperactivity disorder. J. Am. Acad. Child Adolesc. Psychiatry 49, 884–897 (2010).

46. Mick, E. et al. Family-based genome-wide association scan of attention-deficit/hyperactivity disorder. J. Am. Acad. Child Adolesc. Psychiatry 49, 898–905.e3 (2010).

47. Elia, J. et al. Rare structural variants found in attention-deficit hyperactivity disorder are preferentially associated with neurodevelopmental genes. Mol. Psychiatry 15, 637–646 (2010).

48. Lionel, A. C. et al. Rare Copy Number Variation Discovery and Cross-Disorder Comparisons Identify Risk Genes for ADHD. Sci. Transl. Med. 3, 95ra75 (2011).

49. Neale, B. M. et al. Case-control genome-wide association study of attention-deficit/hyperactivity disorder. J. Am. Acad. Child Adolesc. Psychiatry 49, 906–920 (2010).

50. Stergiakouli, E. et al. Investigating the contribution of common genetic variants to the risk and pathogenesis of ADHD. Am. J. Psychiatry 169, 186–194 (2012).

51. Sánchez-Mora, C. et al. Meta-analysis of brain-derived neurotrophic factor p.Val66Met in adult ADHD in four European populations. Am. J. Med. Genet. B. Neuropsychiatr. Genet. 153B, 512–23 (2010).

52. Hinney, A. et al. Genome-wide association study in German patients with attention deficit/hyperactivity disorder. Am. J. Med. Genet. Part B Neuropsychiatr. Genet. 156B, 888–97 (2011).

53. Zayats, T., Johansson, S. & Haavik, J. Expanding the toolbox of ADHD genetics. How can we make sense of parent of origin effects in ADHD and related behavioral phenotypes? Behav. Brain Funct. 11, 33 (2015).

54. Gelernter, J. et al. Genome-wide association study of alcohol dependence:significant findings in African- and European-Americans including novel risk loci. Mol. Psychiatry 19, 41–9 (2014).

55. Gelernter, J. et al. Genome-wide association study of cocaine dependence and related traits: FAM53B identified as a risk gene. Mol. Psychiatry 19, 717–23 (2014).

56. Gelernter, J. et al. Genome-wide association study of opioid dependence: multiple associations mapped to calcium and potassium pathways. Biol. Psychiatry 76, 66–74 (2014).

57. Sherva, R. et al. Genome-wide Association Study of Cannabis Dependence Severity, Novel Risk Variants, and Shared Genetic Risks. JAMA psychiatry (2016). doi: 10.1001/jamapsychiatry.2016.0036

58. Bulik-Sullivan, B. et al. An atlas of genetic correlations across human diseases and traits. Nat. Genet. 47, 1236–1241 (2015).

59. Bulik-Sullivan, B. et al. LD Score regression distinguishes confounding from polygenicity in genome-wide association studies. Nat. Genet. 47, 291–295 (2015).

60. Yang, J., Lee, S. H., Goddard, M. E. & Visscher, P. M. GCTA: a tool for genome-wide complex trait analysis. Am. J. Hum. Genet. 88, 76–82 (2011).

61. Lee, S. H., Yang, J., Goddard, M. E., Visscher, P. M. & Wray, N. R. Estimation of pleiotropy between complex diseases using single-nucleotide polymorphism-derived genomic relationships and restricted maximum likelihood. Bioinformatics 28, 2540–2542 (2012).

62. The International Schizophrenia Consortium. Common polygenic variation contributes to risk of schizophrenia and bipolar disorder. Nature 460, 748–752 (2009).

63. Wray, N. R. et al. Research Review: Polygenic methods and their application to psychiatric traits. J. Child Psychol. Psychiatry 55, 1068–1087 (2014).

64. Ludvigsson, J. F., Otterblad-Olausson, P., Pettersson, B. U. & Ekbom, A. The Swedish personal identity number: possibilities and pitfalls in healthcare and medical research. Eur. J. Epidemiol. 24, 659–667 (2009).

65. Ludvigsson, J. F. et al. Registers of the Swedish total population and their use in medical research. Eur. J. Epidemiol. 31, 125–136 (2016).

66. Ekbom, A. in 215–220 (2011). doi: 10.1007/978-1-59745-423-0_10

67. Ludvigsson, J. F. et al. External review and validation of the Swedish national inpatient register. BMC Public Health 11, 450 (2011).

68. Wettermark, B. et al. The new Swedish Prescribed Drug Register—Opportunities for pharmacoepidemiological research and experience from the first six months. Pharmacoepidemiol. Drug Saf. 16, 726–735 (2007).

